# Structure and origin of the *White Cap* locus and its role in evolution of grain color in maize

**DOI:** 10.1101/082677

**Authors:** Bao-Cai Tan, Jiahn-Chou Guan, Shuo Ding, Shan Wu, Jonathan W. Saunders, Karen E. Koch, Donald R. McCarty

## Abstract

Selection for yellow and white grain types has been central to post-domestication improvement of maize. While genetic control of carotenoid biosynthesis in endosperm is attributed primarily to the *Yellow1* (*Y1*) phytoene synthase gene, less is known about the role of the dominant white endosperm factor *White Cap* (*Wc*). We show that the *Wc* locus contains multiple, tandem copies of a *Carotenoid cleavage dioxygenase 1* (*Ccd1*) gene that encodes a carotenoid-degrading enzyme. A survey of 111 maize inbreds and landraces, together with 21 teosinte accessions reveals that *Wc* is exclusive to maize, where it is prevalent in white-grain (*y1*) varieties. Moreover, *Ccd1* copy number varies extensively among *Wc* alleles (from 1 to 23 copies), and confers a proportional range of *Ccd1* expression in diverse organs. We propose that this dynamic source of quantitative variation in *Ccd1* expression was created in maize shortly after domestication by a two-step, *Tam3L* transposon-mediated process. First, a chromosome segment containing *Ccd1* and several nearby genes duplicated at a position 1.9 Mb proximal to the progenitor *Ccd1r* locus on chromosome 9. Second, a subsequent interaction of *Tam3L* transposons at the new locus created a 28-kb tandem duplication, setting up expansion of *Ccd1* copy number by unequal crossing over. In this way, transposon-mediated variation in copy number at the *Wc* locus created phenotypic variation that provided a foundation for breeding and selection of white grain color in maize.

## Introduction

Structural rearrangements and gene copy-number variation are important components of genetic diversity in plant genomes (Springer et al., 2009; DeBolt, 2010; Hardigan et al., 2016). Although the sources of structural variation are not fully understood, transposons are a potent mechanism of genome remodeling in maize (Fu and Dooner, 2007), including novel genotypes associated with domestication (Studer et al., 2011). In particular, *Ac/Ds* transposons belonging to the *Hat* (*Hobo-Activator-Tam3*) super-family of DNA transposons (Kempken and Windhofer, 2001) have been shown to mediate a rich repertoire of chromosome rearrangements in maize (Ralston et al., 1989; Zhang et al., 1999, 2013, 2014; Huang and Dooner, 2008). The potential of the *Ac/Ds* elements for generating gene duplications, transpositions, deletions and inversions is attributable to three essential features of the *Ac/Ds* system; 1) transposition typically occurs during DNA replication; 2) *Ac/Ds* elements often transpose to sites that are near the donor site in the genome; and 3) compatible ends of nearby elements readily interact to form macro-transposons. Macro-transposition events can produce a range of structural outcomes depending on i) relative orientation of the interacting elements, ii) relative orientation of transposon ends at the insertion site and iii) position of the insertion site and interacting elements relative to nearby replication forks at the time of transposition (Ralston et al., 1989; Zhang et al., 2014; Huang and Dooner, 2008). While, thus far, the *Ac/Ds* system has been studied extensively in maize, the broad distribution of *Hat* transposons in plant as well as other eukaryotic genomes (Kempken and Windhofer, 2001) indicates that the mechanisms demonstrated maybe a widespread source of structural variation.

Carotenoid content of the endosperm is a key trait that affects both the nutritional and aesthetic qualities of the maize grain (Brink, 1930; Buckner et al., 1990, 1996, add Tracy?). Both yellow- and white-endosperm varieties are agronomically important (Poneleit, 2001). The white endosperm typical of teosinte grain is presumed to be the ancestral phenotype (Palaisa et al., 2004). In maize, carotenoid biosynthesis in endosperm requires dominant alleles of the *Y1* gene that confer expression of phytoene synthase in both seed and plant tissues. In contrast, recessive *y1* alleles have low phytoene synthase in endosperm, and produce a white grain phenotype (Buckner et al., 1996). Association-genetic studies indicate that human selection for a dominant *Y1* allele occurred during domestication of yellow grain maize (Buckner et al., 1990; Palaisa et al., 2003, 2004; Zhu et al., 2008). However, at least two genes determine white versus yellow endosperm. In addition to the recessive *y1*, a white-endosperm can result from action of dominant alleles of *White cap (Wc).* The dominant nature of *Wc*, which has been known for at least a century (White, 1917; Brink, 1930), implies a mechanism for negative regulation of carotenoid accumulation in the developing maize kernel. *Wc* occurs in commercially important white- and sweet-corn varieties (Hannah and McCarty, 1991).

The Carotenoid Cleavage Dioxygenase 1 (CCD1) enzyme is a broad-specificity 9,10 (9’, 10’) carotenoid dioxygenase that catalyzes cleavage of diverse carotenoids to their corresponding apo-carotenoid products (Tan et al., 1997, 2003; Vogel et al., 2008; Sun et al., 2008). In animals, apo-carotenoids such as retinoids and vitamin A are derived from specific cleavage of carotenoids (Schwartz et al., 1997) and serve as signaling molecules in animals (Giguere et al, 1987). In plants, apo-carotenoids are precursors for two important hormones, abscisic acid (ABA) and strigolactone (Tan et al., 1997; Schwartz et al., 1997; Gomez-Roldan et al., 2008; Umehara et al., 2008; Zeevaart et al., 1989).

Here we show that the *Wc* locus, which confers a white-endosperm phenotype, contains multiple tandem copies of the *Ccd1* gene. Alleles of W*c* can have between 1 and 23 copies of a 28-kb tandem repeat that contains *Ccd1* and two downstream genes, glutamyl tRNA acyl transferase *(Tglu)* and a cytochrome P450 *(P450).* Moreover, we find that *Ccd1* mRNA levels in diverse tissues of *Wc* inbreds vary in direct proportion to *Ccd1* copy number. Our analyses of *Wc* structure and distribution based on bac- and whole-genome-sequence (wgs) data indicate that the *Wc* locus was created by separate macro-transposition and gene amplification events – both mediated by interactions between closely-spaced *Tam3-like (Tam3L)* transposons. First, a pair of *Tam3L* elements formed a macro-transposon that created a duplication of the chromosome segment containing *Ccd1* and several nearby genes at a position 1.9 Mb proximal to the progenitor *Ccd1* locus *(Ccd1r).* Next, a subsequent interaction of *Tam3L* transposons at the new locus formed a 28-kb tandem duplication that initiated further expansion of repeat copy-number by un-equal crossing-over. Although the *Wc* phenotype is most dramatic in the endosperm of yellow-grained, *Y1* genotypes, *Wc* occurs most often in *y1* varieties that lack capacity for carotenoid biosynthesis in endosperm. We suggest that *Wc* intensifies the white-grain phenotype of recessive y1, thus providing a basis for human selection of the *Wc y1* genotype. Our results indicate that variation in *Ccd1* copy-number and expression due to *Wc* enriched the genetic foundation for breeding and selection for grain color during post-domestication improvement of maize.

## Results

### *Wc* contains a *Ccd1* gene cluster that confers high *Ccd1* expression

In a *Y1* genetic background, which leads to biosynthesis of yellow carotenoids in the endosperm, the dominant *Wc* allele confers a dosage-dependent white endosperm phenotype (Fig 1a). In the triploid endosperm, kernels that have a single dose of *Wc* have a pale-yellow or white crown, reflecting a partial inhibition of carotenoid accumulation. In contrast, kernels with three doses of *Wc* have a nearly white endosperm. A gradation of yellow to white kernel phenotypes can thus be discerned on a self-pollinated ear of a *Wc wc Y1 Y1* heterozygote consistent with the four expected gene-dosage classes (Fig 1a).

**Figure 1.**
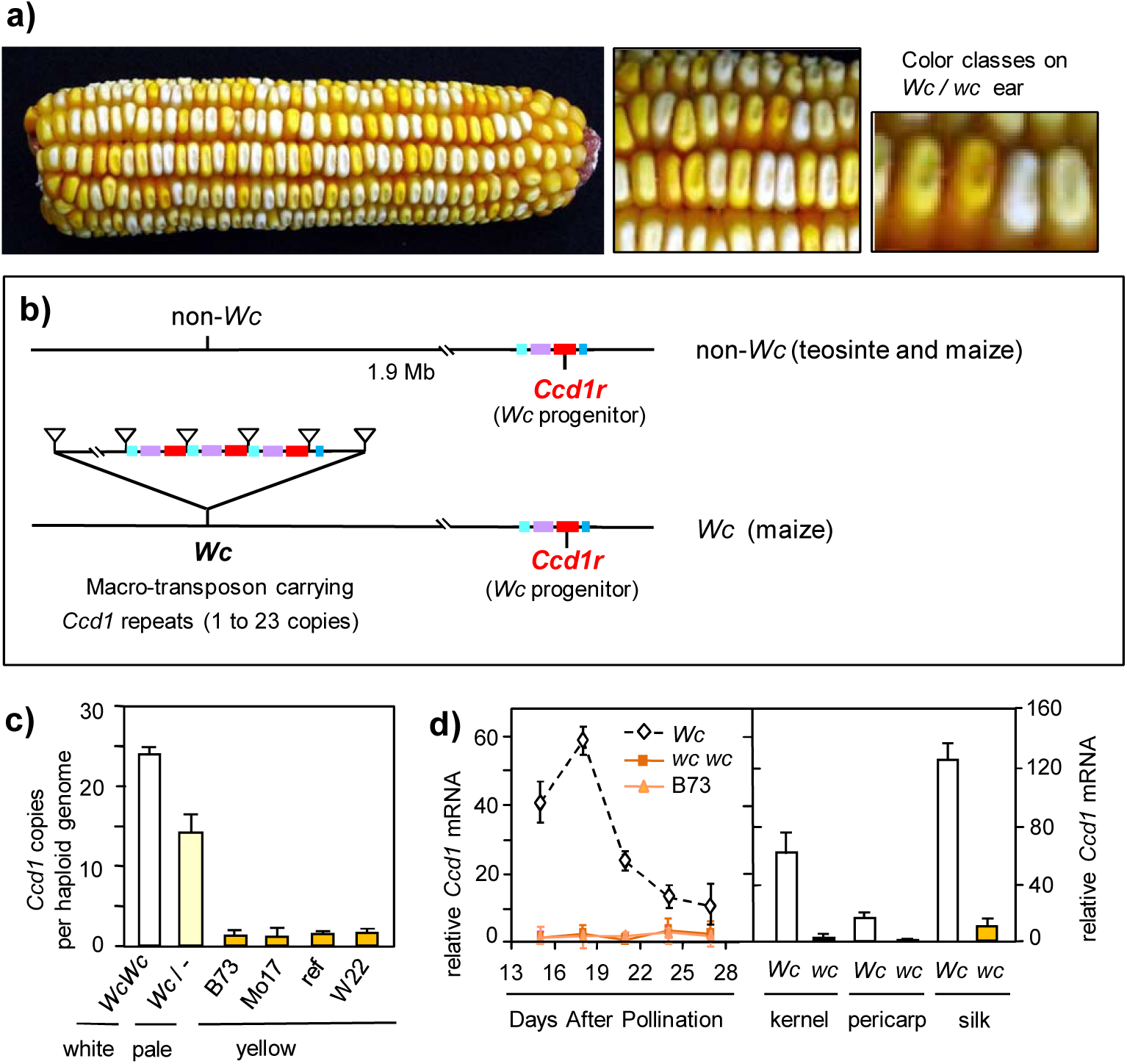
*White Cap* (*Wc*), a dominant inhibitor of carotenoid accumulation in maize endosperm encodes a macro-transposon that contains 1 to 23 tandem copies of *Ccd1* (*Carotenoid* c*leavage dioxygenase-1*) causing elevated expression in diverse tissues. (**a**). Kernels on a self-pollinated ear of a *Wc wc* heterozygote in a *Y1 Y1* background segregate four classes of yellow pigmentation distinguished by *Wc* dosage in the triploid endosperm. Kernels with three doses of *Wc* (25% of kernels) have a fully-white phenotype whereas 0 doses (25% of kernels) conditions a normal yellow phenotype. Intermediate *Wc* dosage classes have pale yellow phenotypes. (**b**). Diagram comparing *Wc* (maize) and *non-Wc* (maize and teosinte) chromosomes. The *Ccd1r* reference locus is presumed to be present in all maize and teosinte haplotypes (see Table 1). In *Wc* maize a macro-transposon insertion carries from 1 to 23 tandem copies of the *Ccd1* gene is located 1.9 Mb proximal to *Ccd1r*. The transposed chromosome segment includes *Ccd1* (red), and neighboring genes *Tglu* (purple), *P450* (light turquoise), and *Rpl21* (blue). Supporting evidence is described herein (see also subsequent figures). (**c**). Quantification of *Ccd1* copies per genome in the Wc reference allele (Wc Wc), a Wc heterozygote from a SilverQueen B73 F1 hybrid, and several non-Wc (wc) inbreds (ref, isogenic non-Wc control inbred). (**d**). Expression of *Ccd1* during seed development. mRNA abundance per ng total RNA determined by qPCR is expressed in relative units (see Materials and Methods).

Fig 1b summarizes the structure of the *Wc* locus and its relationship to the *Ccd1 reference* (*Ccd1r)* locus in *Wc* and non-Wc (*wc*) haplotypes. Evidence presented below indicates that *Wc* is derived from insertion of a macro-transposon that duplicated a region including *Ccd1* and several nearby genes at a position 1.9 Mb proximal to the *Ccd1r* locus on the long-arm of chromosome 9. Southern blot analysis (Supplemental Fig. S1a) showed that *Wc* co-segregated with a gene cluster that includes multiple copies of the *Ccd1* coding sequence. BamHI restriction digests probed with a *Ccd1r* cDNA yield a single, intense, 6.1-kb band, indicating that the multiple copies are highly homogeneous. Digests with additional enzymes indicate that the *Ccd1* copies are embedded in a homogeneous repeat at least 16 kb long (Supplemental Fig S1b.). In line with these results, our *Ccd1* cDNA probe detects an intense FISH signal on the long-arm of chromosome 9 in *Wc* plants (Han et al., 2007).

To quantify *Ccd1* copy number in *Wc* and non-Wc (*wc*) genotypes (Fig 1c), we developed a gene-specific, real-time PCR assay for *Ccd1* utilizing the well-characterized *Vp14* gene (Tan et al., 1997) as a single-copy internal control. Consistent with the Southern blot results, PCR data indicate that genomes of B73 and other yellow inbreds carry a single-copy of *Ccd1*, which we attribute to *Ccd1r*. In contrast, plants homozygous for the *Wc* reference allele are estimated to have 23.9 (+/- 2.0) copies per genome. In agreement with this estimate, heterozygotes carrying a *Wc* allele extracted from “Silver Queen” sweet corn (Hannah and McCarty, 1991) have an estimated 13.5 (+/-2.4) copies. This is an expected value for a heterozygote carrying 24 copies on one chromosome and 1 copy on the other [(24+1)/2 = 12.5 copies per genome].

Based on these results, we reasoned that amplification of *Ccd1* copy number at the *Wc* locus could account for the dominant white phenotype by elevating expression of the broad-specificity CCD1 9,10 carotenoid dioxygenase activity (Vogel et al., 2008) in the endosperm. As shown in Fig 1d, relative expression of *Ccd1* mRNA is indeed markedly higher throughout development of *Wc* kernels compared to isogenic non-Wc and B73 inbreds. In *Wc* kernels, *Ccd1* expression peaks during mid-grain fill at 18 DAP then declines gradually toward maturity. In addition, *Wc* causes elevated *Ccd1* expression in diverse tissues including silks and pericarp.

### Structure of the *Wc* locus

To determine the structure of the *Wc* locus, we constructed a bac library from the *Wc* reference stock and isolated a 106-kb bac clone that spanned one boundary of the *Ccd1* gene cluster. The bac sequence assembly diagrammed in Fig 2a was confirmed by 1) restriction mapping, 2) sequencing of bac-ends and selected subclones, 3) genomic Southern blot data (Supplemental Fig S2a-b), 4) consistency with whole-genome-sequence (wgs) data from diverse *Wc* inbreds (Table 1), and 5) by mapping and partial sequencing of a second overlapping bac clone (data not shown). To aid interpretation of the *Wc* assembly, the bac sequence was then aligned to the region surrounding the single-copy *Ccd1r* gene in the B73 reference genome (Fig 2b-c). In addition, we cloned and sequenced a 12-kb region containing the *Ccd1r* gene from the teosinte (*Zea mays* spp. *parviglumis*) genome (Fig 2c, Supplemental Fig S2b). The 106-kb *Wc* bac sequence includes 3 tandem copies of *Ccd1* arranged in direct orientation. Each *Ccd1* copy is embedded in a 28-kb direct repeat that also includes a glu-tRNA acyltransferase (*Tglu*, GRMZM2G057491) and a cytochrome P450 (GRMZM2G057514) gene, *P450*. The *Tglu* and *P450* genes are located downstream of *Ccd1r* in the B73 reference genome (Fig 2b-c). The rightmost *Ccd1* copy in the bac sequence contains a 4,460-bp, *Tam3-like* transposon insertion (*Tam3Lb*) near its 5’-prime end. As expected, *Tam3Lb* is flanked by an 8-bp, host-site duplication typical of the *Hat* transposon family. Although multiple *Tam3L* copies are detected in the B73 genome (data not shown), the *Tam3L* transposon family has not previously been characterized in maize.

**Table 1.**
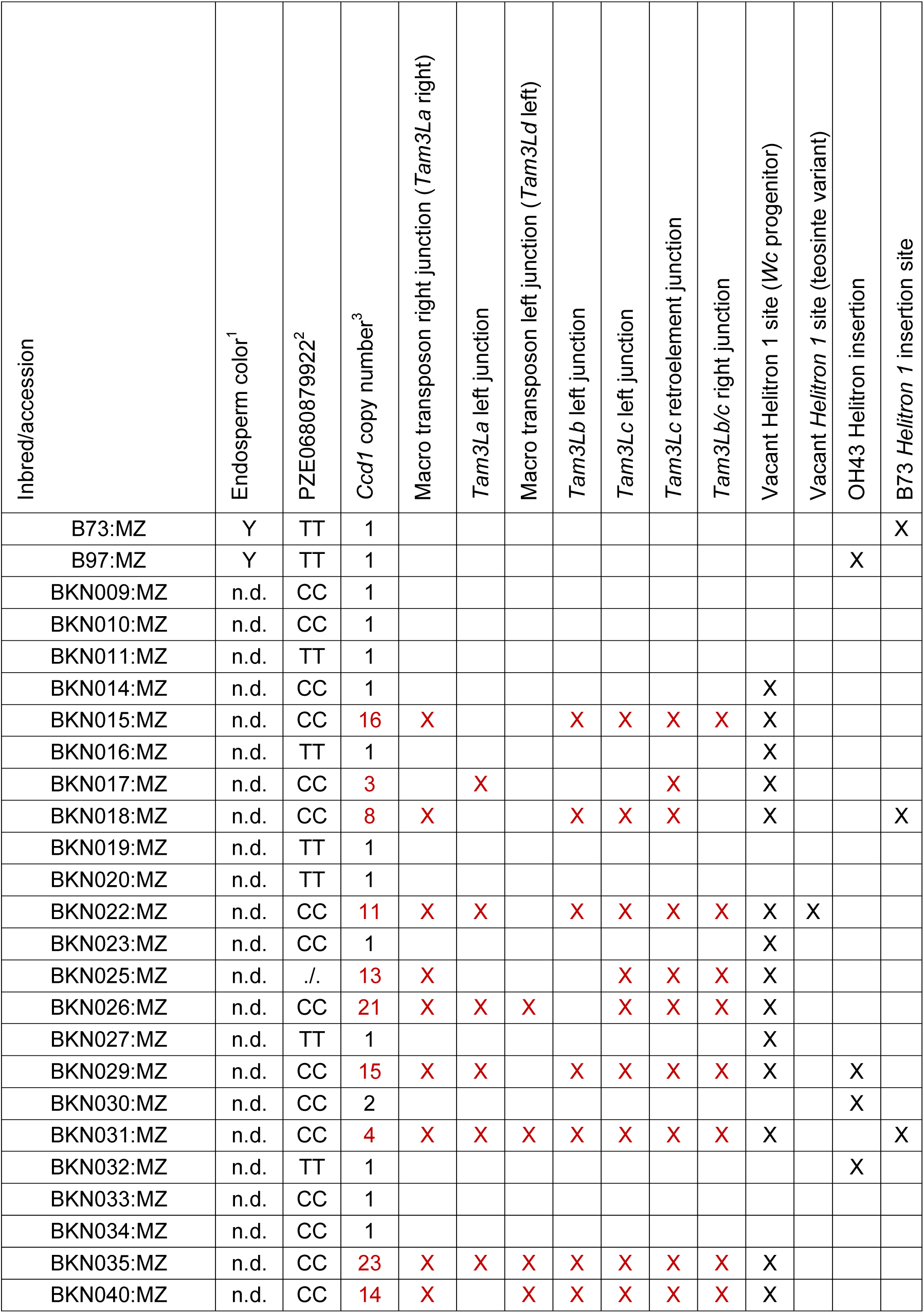
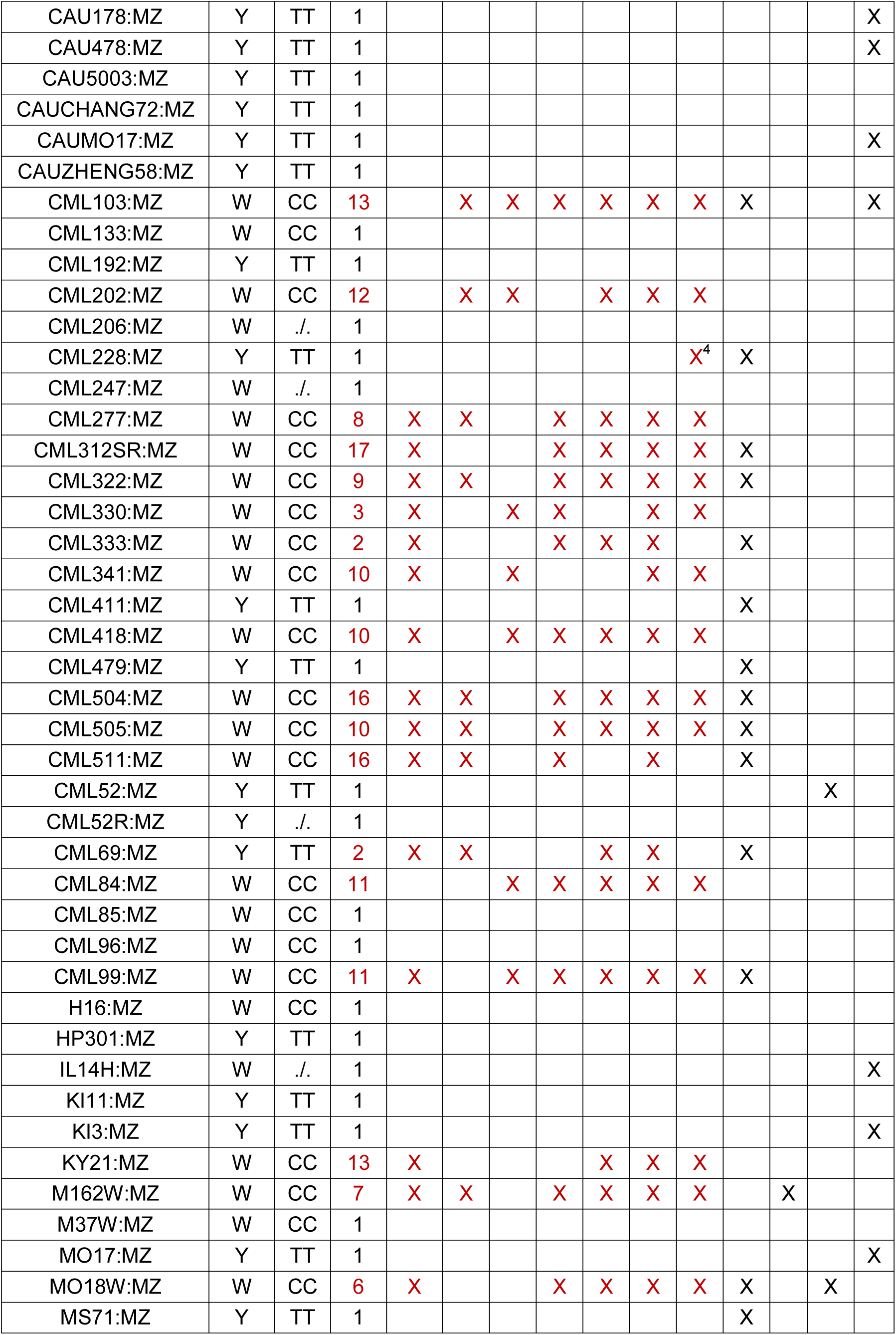
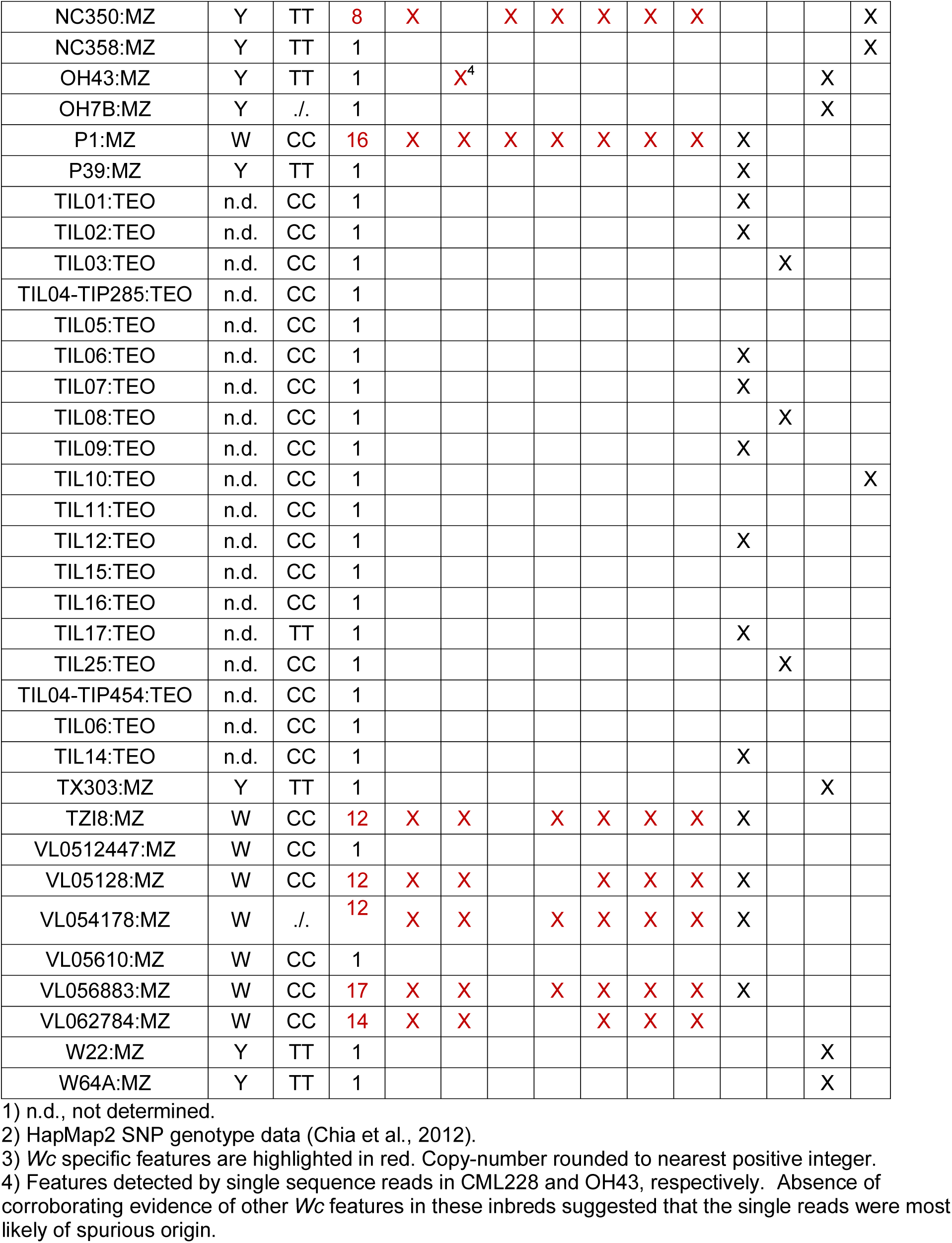
Sequence features detected in genomes of maize and teosinte accessions

**Figure 2.**
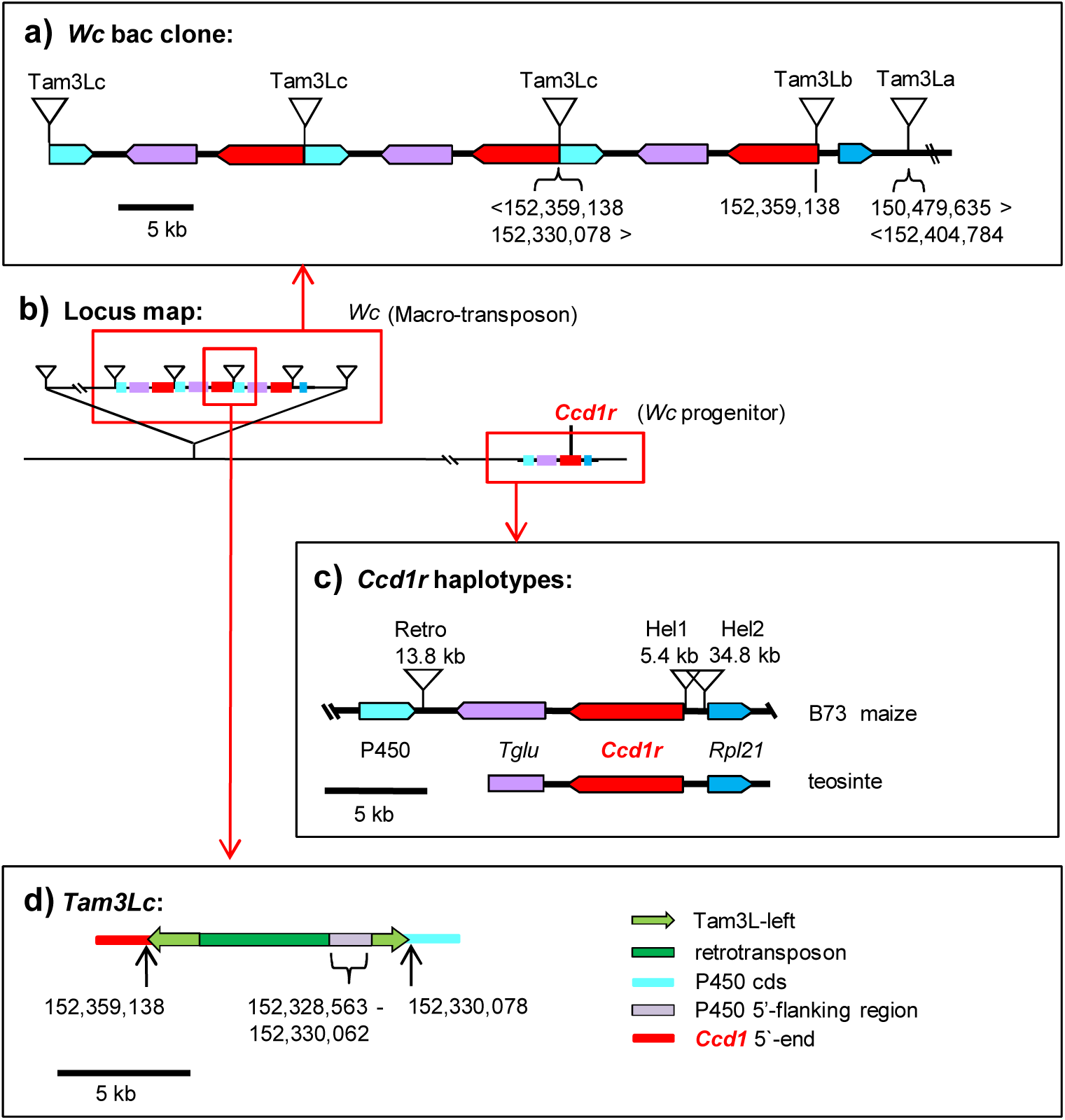
Structure of the *Wc locus and Ccd1r haplotypes of maize and teosinte.* (**a**) Sequencing and assembly of a bac clone containing one border of the *Wc* locus revealed two complete and one partial copy of a 28 Kb repeat that included copies of *Ccd1* (red) and nearby *Tglu* (purple) and *P450* (turquoise) genes. The repeats are punctuated by a composite *Tam3L* like sequence (*Tam3Lc*), which is shown in greater detail in part **d**. The rightmost copy of *Ccd1* is bordered by an intact *Tam3L* transposon (*Tam3Lb*) with a flanking 8 bp host site duplication. The 10 kb segment between *Tam3Lb* and a second *Tam3L* transposon (*Tam3La*) is co-linear with the region upstream of *Ccd1r* in the B73 reference and teosinte genomes that includes *Rpl21* (blue, see part C). The 13.5 kb sequence that extends from *Tam3La* to the right end of the bac clone (abridged for clarity) aligns to sequence located 1.9 Mb proximal to *Ccd1r* in the B73 reference. Consequently, *Tam3La* is not flanked by an 8-bp host site duplication. (**b**) Location of Wc relative to the *Ccd1r* progenitor locus on the long-arm of chromosome 9 (see Fig 1B). **(c)** Comparison of *Ccd1r* haplotypes found in maize (B73 reference) and teosinte (data on distributions of *Ccd1r* haplotypes in maize and teosinte are summarized in Table 1). The teosinte structure was determined by sequencing a 12kb region containing *Ccd1r* in Z. m. parviglumis. (**d)**. Structure of *Tam3Lc*, a composite transposon-like element that punctuates the *Wc* 28 kb repeats. Numbers in parts **a** and **d** indicate coordinates of sequence alignments to the B73 genome (version 3).

### *Wc* contains a *Tam3L* macro-transposon

In the bac sequence, a 6,573-bp region flanking the right side of the *Tam3Lb* transposon is co-linear with genomic sequence upstream of *Ccd1r* in the teosinte genome (Fig 2c) and includes a copy of the neighboring gene for a *Ribosomal-large-subunit-protein-21* (*Rpl21*, GRMZM2G089421). This region of co-linearity with the ancestral *Ccd1r* haplotype is bordered on the right by a second *Tam3L* element (*Tam3La*). Although the *Tam3La* transposon is intact, two features of its flanking sequences indicated that *Tam3La* likely forms one boundary of a macro-transposon. First, *Tam3La* is not flanked by an 8-bp direct duplication. Second, sequences on the left and right sides of *Tam3La* are not contiguous in the B73 reference genome. Instead, sequence to the right of *Tam3La* aligns to a position 1.9 Mb proximal to the *Ccd1r* locus in the B73 genome.

On this basis, we hypothesized that the other boundary of the *Wc* locus would be delimited by another *Tam3L* transposon (*Tam3Ld*). We further anticipated that *Tam3Ld* would have a left flanking sequence that 1) was contiguous in the reference genome with sequence flanking the right side of *Tam3La* (Fig 2a), and 2) shared a matching 8-bp host site duplication with *Tam3La*. As hypothesized, we were indeed able to identify the predicted *Tam3Ld* junction sequence that defines the left boundary of the *Wc* macro-transposon in genomic DNA of *Wc* plants. We amplified the *Tam3Ld* left junction using a combination of TAIL-PCR and PCR with primers specific for the left arm of *Tam3L* and the expected flanking sequence (Fig 3). In addition, we detected the *Tam3Ld*-left junction sequence in wgs data from multiple *Wc* inbreds (Table 1).

**Figure 3.**
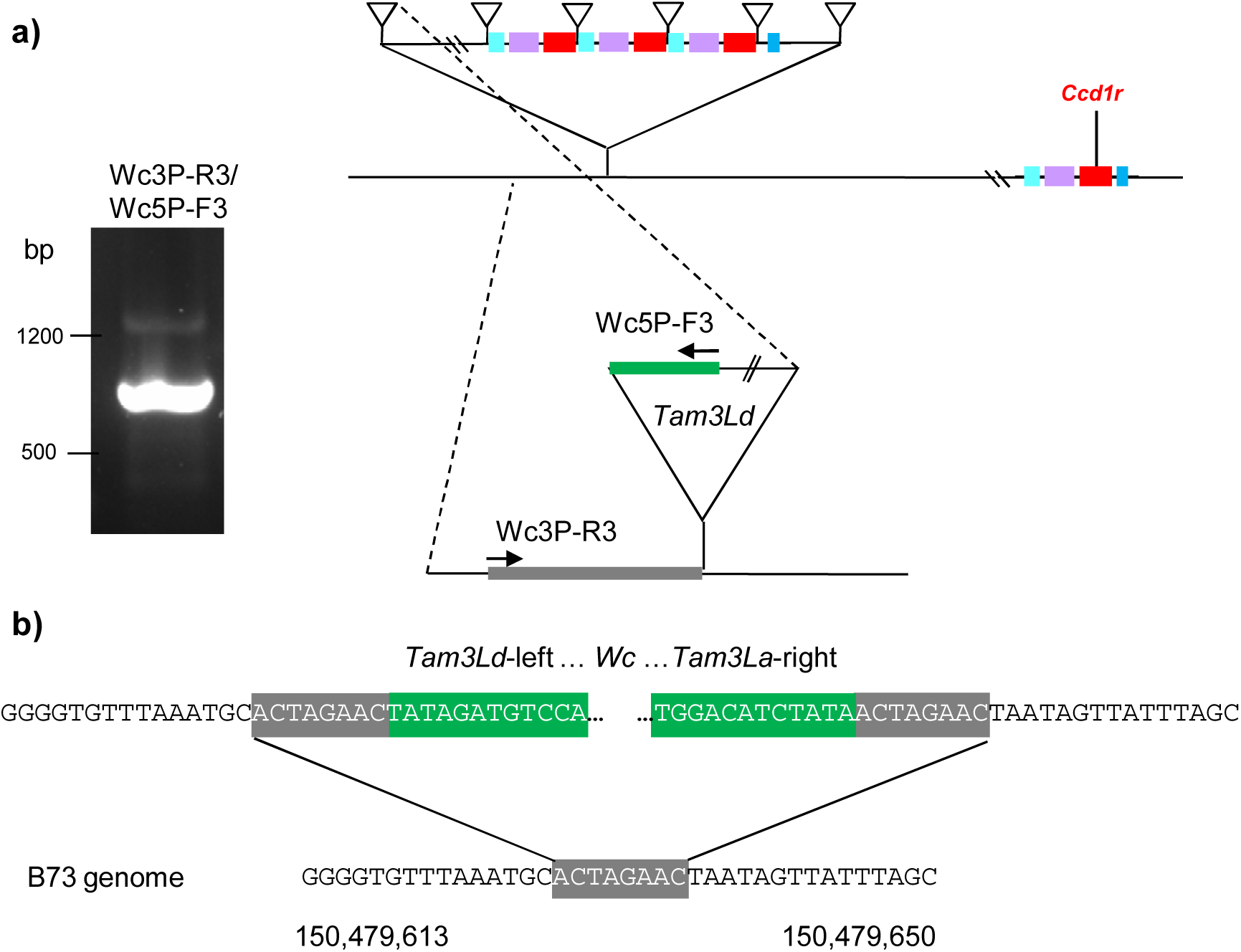
Identification of the T*am3Ld* insertion marking the proximal border of the *Wc* m acro-tr ansposon. (**a**) PCR am pl if ication of the *Tam3Ld* lef t j uncti on f rom *Wc* pl ant s. Candidat e *Tam3L*-l ef t j unct i on sequences were amplified by TAIL-PCR anchored by locus specific primers in the predicted location of *Tam3Ld* (see Materials and Methods). The TAIL-PCR sequences were used to design a pair of sequence specific primers, Wc3P-R3 and Wc3P-F3, that amplified a 794 bp junction fragment (agarose gel electrophoresis image). The locations of sequence-specific primers are diagramed on the right. Green indicates transposon sequence. The 8bp target site duplication is grey. (**b**) Confirmation of the *Wc* macro-transposon insertion site. The*Tam3Ld* junction sequence aligned to the B73 reference genome at a position that was contiguous with the right flanking sequence of Tam3La and included a matching 8bp target site duplication (grey). The *Tam3Ld* and *Tam3La*, 12 base inverted terminal repeats are colored green. B73 chromosome 9 genome coordinates (version 3) are indicated below the sequence.

### *Wc* repeats are punctuated by a composite *Tam3L* sequence

Each 28-kb repeat is bordered by a 9,980-bp transposon-like sequence (*Tam3Lc*) that has two *Tam3L*-left termini (Fig 2d, Supplemental Fig S3) with two unrelated sequence fragments sandwiched between them. The two internal sequences include a 4,835-bp fragment of an uncharacterized retrotransposon and a 1,508-bp segment that aligns to genomic sequence flanking the 5’-end of the *P450* gene in the B73 reference genome. Sequence derived from the *P450* flanking region is nearly contiguous with the right junction of *Tam3Lc*, except for a deletion of 16 bp at the insertion site. These features are consistent with an abortive transposition that inserted a single *Tam3L* end upstream of *P450.* The right end of *Tam3Lc* has several polymorphisms in common with *Tam3La*-left (Supplemental FigS2) that distinguish it from *Tam3Lb-*left, whereas the opposite end of *Tam3Lc* is derived from the left terminus of *Tam3Lb*.

### Creation of *Wc* by macro-transposition and gene amplification

The structural features described above indicated that the *Wc* locus was created by a series of interactions of between *Tam3L* elements that 1) initially transposed a copy of the *Ccd1r* locus to a position 1.9 Mb upstream of the progenitor locus, and 2) initiated amplification of a 28-kb segment of the transposed copy. A proposed series of events that account for the structure depicted in Fig 2, is outlined in Supplemental Fig. S4. In this scenario, creation of the *Wc* locus was preceded by a series of *Tam3L* insertions in the ancestral *Ccd1r* locus. This resulted in a pair of *Tam3L* elements that flanked the region containing *Ccd1r* and neighboring *Rpl21, Tglu* and *P450* genes. As a replication fork moved through this segment *Tam3Ld*-left and *Tam3La*-right then formed a macro-transposon that inserted at an un-replicated site 1.9 Mb proximal to the *Ccd1r* locus. We suggest that a subsequent interaction of *Tam3L* elements at the new locus duplicated a segment containing *Ccd1*, thus enabling expansion of repeat-number by unequal crossing-over. In some configurations *Ac* is capable of creating tandem duplications by initiating rolling-circle-replication or re-replication (Ralston et al., 1989; Zhang et al., 2014). In instances where *Ac* has induced re-replication of DNA, Zhang et al. (2014) observed that stalling of the replication fork prevented extensive rolling-circle-replication. While a similar mechanism may have created the initial tandem repeat in *Wc*, the precise origin of the composite *Tam3Lc* element is unclear. The incorporation of an interior sequence derived from the upstream flanking region of the *P450* gene suggests that *Tam3Lc* was formed in part by an abortive transposition that inserted one arm of a transposon - possibly derived from *Tam3La* - upstream of *P450*. One speculative possibility is that a macro-transposition involving *Tam3La*-left and *Tam3Lb*-right aborted during replication, resulting in a fractured chromosome with one or more double stranded breaks. Formation of *Tam3Lc* by repair reactions that fused nearby *Tam3La*-left and *Tam3Lb*-left fragments could plausibly have created a circular template that initiated transient re-replication of *Ccd1*.

### Independent diversification of *Ccd1r* and the *Wc* locus

Subsequent to creation of *Wc*, the *Ccd1r* progenitor and *Wc* continued along separate paths of evolution and diversification (Fig 2c). Comparison of the *Ccd1r* sequences from B73 and ancestral teosinte revealed that the B73 haplotype contains helitron transposon insertions near the transcription start sites of both the *Ccd1r* and *Rpl21* genes. The helitron insertions displaced upstream flanking sequences of both genes, potentially altering their regulation. A search of wgs data detected the B73 *Hel1* junction sequence in at least 13 of 84 maize inbreds and 1 of 19 teosinte accessions (Table 1). At least 4 *Wc* inbreds also carried *Hel1* (B73 type) *Ccd1r* alleles. As expected based on the *Wc* bac sequence, the *Ccd1* promoter sequence lacking the *Hel1* insertion was found in a majority of inbreds that carry *Wc*. This sequence also occurred in at least 8 non-*Wc* maize inbreds and 8 of 19 teosinte accessions indicating that *Ccd1r* alleles resembling the *Wc* progenitor exist in both maize and teosinte (Table 1). Evidence of a second, independent helitron insertion located in a similar position upstream of *Ccd1r* was detected in OH43 (data not shown). The OH43 promoter variant was detected in at least 11 of 84 maize accessions, but in none of the teosinte accessions (Table 1).

### *Wc* alleles account for extensive *Ccd1* copy-number variation in maize

To evaluate *Ccd1* copy number variation in maize, we analyzed wgs data from 101 maize and teosinte genomes represented in the HapMap2 resource (Chia et al., 2012). The number of *Ccd1* copies was estimated by k-mer frequency analysis of raw wgs data from the HapMap2 collection using JELLYFISH (Marçais and Kingsford, 2011). *Ccd1* copy-number per genome was estimated by determining the frequencies of *Ccd1*-specific 22-mers in each data set. The counts of *Ccd1* 22-mers in wgs data were normalized to a set of control 22-mers. The control set consisted of 124M, single-copy 22-mers derived from the B73 filtered gene set (gramene.org). The estimated effective depth of coverage obtained for each genome is summarized in Supplemental Table S1. Of the 84 maize genomes represented in the HapMap2 collection, 36 (42%) had an estimated *Ccd1* copy number of 2 or greater (Table 1, rounding to nearest integer). The highest copy number detected, 23 copies per genome in landrace accession BKN035, was comparable to the qPCR-based estimate of 24 copies in the *Wc* reference stock (Fig 1c). All accessions contained at least one *Ccd1* copy which we attributed to the *Ccd1r* locus. In contrast, with one exception, all maize genomes that had 2 or more *Ccd1* copies were confirmed to also contain a suite of sequence features that are specific to the *Wc* locus (Table 1). The exception, landrace BKN030, having an estimate of 2 *Ccd1* copies, contained no evidence of *Wc-*specific features. Therefore, in nearly all cases examined, presence of additional *Ccd1* copies in maize inbreds could be attributed to the *Wc* locus.

### *Wc* was not detected in teosinte

Each of the 19 teosinte genomes represented in the wgs collection show 22-mer frequencies indicating a single copy of *Ccd1* at the progenitor *Ccd1r* locus (Table 1). Moreover, no *Wc-*specific sequence features were detected in teosinte genomes. In line with these data, Southern-blot analysis detected a single *Ccd1* copy in two *Z. m. parviglumis* accessions and one accession of *Z. m. Mexicana* (Supplemental Fig S2). Together these results suggest that the *Wc* locus is unique to maize.

### Structural heterogeneity in *Wc* alleles

Our model for the *Wc* locus predicts that *Ccd1* and *Tglu* copy number should vary among *Wc* alleles in a constant ratio. We found that *Ccd1* and *Tglu* copies were indeed highly correlated (R^2^=0.98; Fig. 4a), though the average ratio of *Tglu* to *Ccd1* copies was somewhat less than the expected 1:1 ratio (0.82:1). This could be explained if the initial tandem duplication depicted in Fig. S4d ended between the *Ccd1* and *Tglu* genes. On that basis, subtracting one copy of *Ccd1*, gives an average ratio in *Wc* inbreds of 1.06 ±0.05 [i.e. *Tglu* copies/(*Ccd1* copies −1) = 1.06] as predicted if the terminal repeat is truncated.

**Figure 4.**
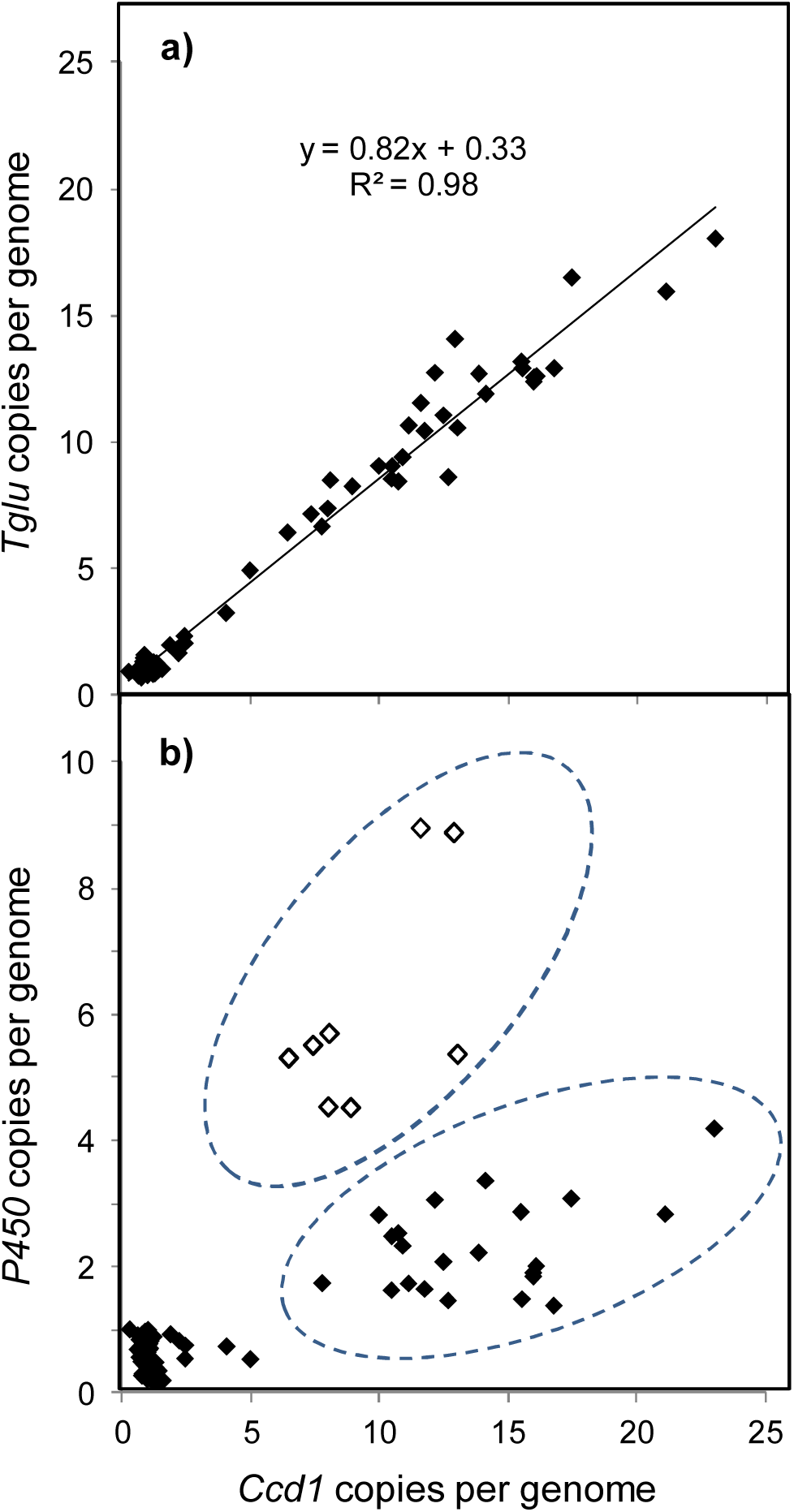
Structural heterogeneity in *Wc* repeats. (**a**) *Ccd1* and *Tglu* copy number vary uniformly in *Wc* alleles. *Ccd1* and *Tglu* copy numbers were estimated using k-mer frequency analysis. The line was determined by linear regression with the equation and R^2^ value indicated. (**b**) Variation in *P450* copy number relative to *Ccd1* reveals structural heterogeneity in *Wc* repeats. Gene copy number was estimated as described in Materials and Methods. Two groups of *Wc* alleles that differ in *P450:Ccd1* copy number ratio are indicated by the dashed ovals. Open symbols, alleles having comparatively high *P450:Ccd1* ratios.

In contrast to the uniform *Tglu:Ccd1* copy-number ratio, the *P450:Ccd1* ratio varied among *Wc* alleles, indicating structural heterogeneity in the *Wc* repeats (Fig 4b). We postulate that alleles with higher *P450:Ccd1* ratios (open symbols in Fig 4b) predominantly contain repeats with the canonical structure (as delineated in Fig. 2a), whereas alleles with lower *P450:Ccd1* ratios include a subset of repeats that have lost most or all of the *P450* sequence. The exact structure of the truncated repeat could not be determined from these data.

### *Ccd1* expression is directly proportional to *Ccd1* copy number

To determine whether the extensive copy number variation at *Wc* is correlated with gene expression, we analyzed RNAseq data from the 27 diverse NAM inbreds (Yu et al., 2008) available at the QTELLER.org database. Based on our analysis (Table 1), 10 inbreds in the NAM collection carry *Wc* alleles with *Ccd1* copy numbers ranging from 2 to 12 copies per genome. As shown in Fig 5, *Ccd1* mRNA levels are highly correlated with *Ccd1* copy number in root, ear, tassel, shoot and shoot apex transcriptomes revealing a nearly linear relationship in diverse tissues (R^2^ values ranging from 0.79 to 0.88 within tissues; R^2^ =0.94 for relative expression normalized over all tissues). By contrast, expression of the adjacent *Tglu* gene shows no discernible relationship to gene dosage despite having a copy-number range comparable to that of *Ccd1* (R^2^=0.05 for relative expression overall, data not shown).

**Figure 5.**
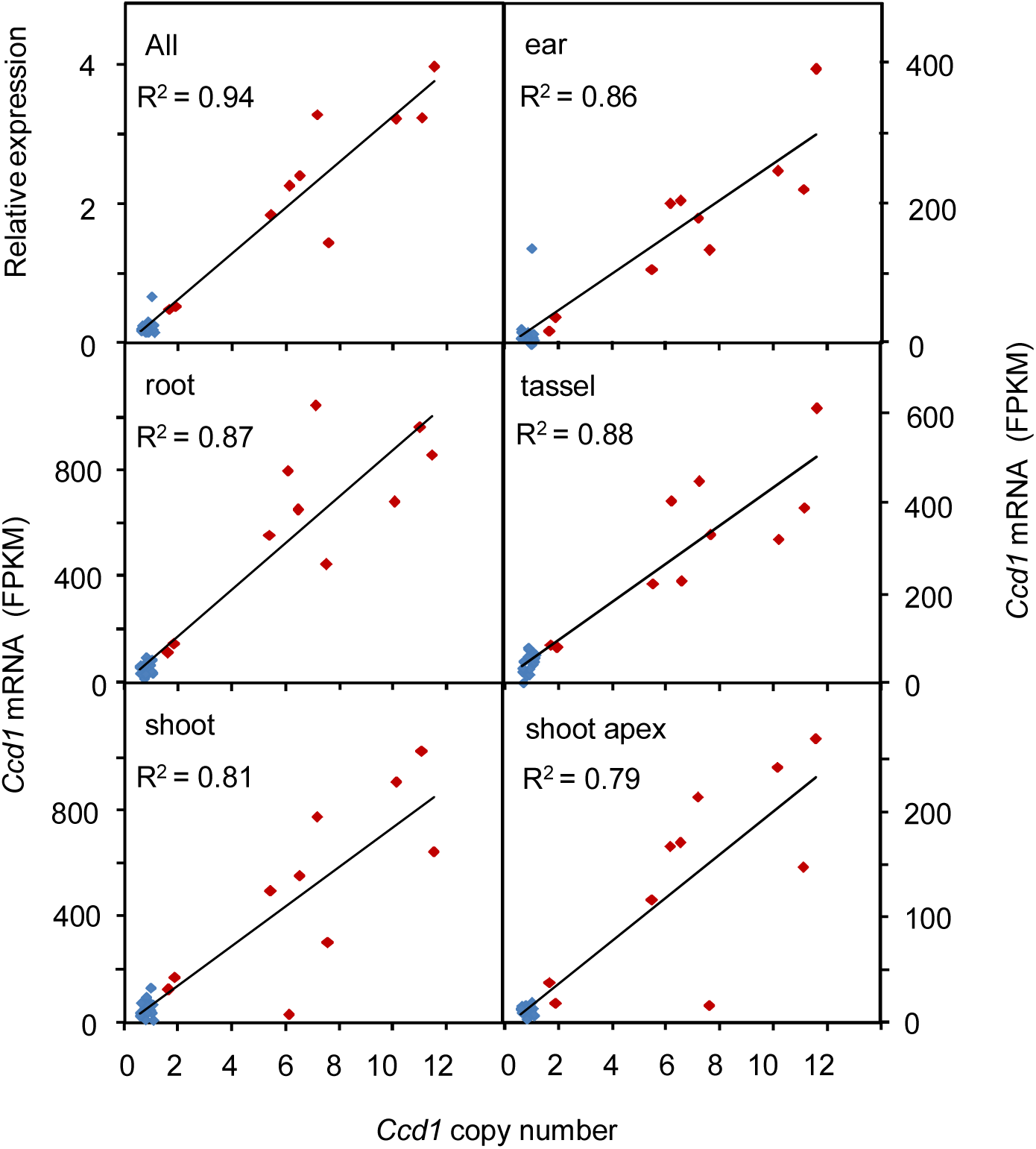
Gene expression is proportional to *Ccd1* copy number. The relationship between *Ccd1* copy number and gene expression in five tissues (root, shoot, ear, tassel, and shoot apex as indicated) was determined for the 27 inbred parents of the NAM population (Yu et al., 2008) using RNAseq data obtained from QTELLER.org. The upper left plot shows the correlation of copy number with relative expression averaged over all tissues using values normalized to the means of each tissue. Red diamonds, inbreds that have the *Wc* locus; blue diamonds, non-Wc inbreds. Lines and R^2^ values were determined by linear regression.

### *Wc* is often associated with recessive *y1* in white-grain maize

The presence of *Wc* in ‘*Silver Queen*’ sweetcorn (Fig 1), inbred A188 (Supplemental Fig S2) and other white inbreds (Stinard, 2010) indicates that *Wc* often occurs in modern inbreds especially those that also carry recessive *y1* alleles. Although *a priori Wc* would not be expected to have an obvious phenotype in *y1* endosperm due to a low capacity for synthesis of CCD1 substrates, there is evidence that *Wc* enhances the white endosperm phenotype of *y1* in backgrounds that carry dominant alleles of *Brown-aleurone-1* (*Bn1*) [Stinard, 2010; Supplemental Fig S6). A survey of inbreds and landraces from diverse geographical locations using Southern blots (Table 2) confirms that *Wc* is present in a majority of white accessions (16 of 25, 64%), but not limited to white maize. At least 6 yellow or mixed-color landraces from Central and South America contain *Wc* alleles. Grain-color phenotype data were available for an additional 60 maize inbreds in the HapMap2 collection (Fig 6). Consistent with the survey results in Table 2, at least 23 of 33 white-grain accessions (70%) contain *Wc* alleles, whereas only 2 of 27 yellow-grain inbreds carry *Wc*. The two exceptions are CML69 (2 *Ccd1* copies) and NC350 (8 *Ccd1* copies). In addition, we noted that within this set of 60 inbreds, white- and yellow-grain color phenotypes, respectively, correlate with C and T variants of SNP PZE0680879922 (Chia et al., 2012). PZE0680879922 is located upstream of the *y1* gene, indicating that it could be used as a marker for the *y1* genotype. On that basis, we infer that as many as 31 of 35 *Wc* accessions (89%) in the HapMap2 collection carry both *Wc* and a recessive *y1* allele.

**Table 2.**
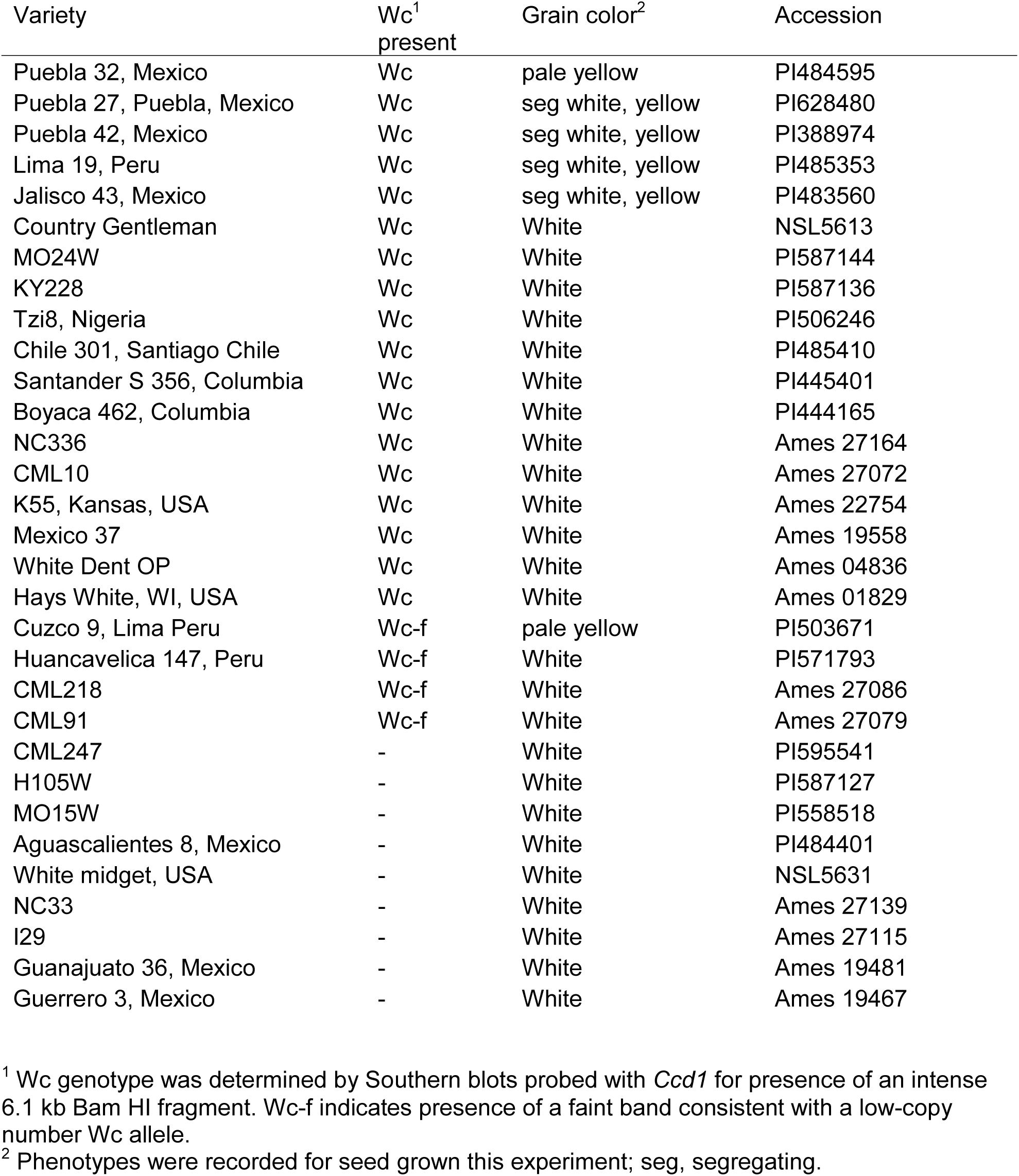
Distribution of *Wc* in white maize inbreds and landraces

| Variety | *Wc*<sup>1</sup><br>present | Grain color<sup>2</sup> | Accession |
| --- | --- | --- | --- |
| Puebla 32, Mexico | *Wc* | pale yellow | PI484595 |
| Puebla 27, Puebla, Mexico | *Wc* | seg white, yellow | PI628480 |
| Puebla 42, Mexico | *Wc* | seg white, yellow | PI388974 |
| Lima 19, Peru | *Wc* | seg white, yellow | PI485353 |
| Jalisco 43, Mexico | *Wc* | seg white, yellow | PI483560 |
| Country Gentleman | *Wc* | White | NSL5613 |
| MO24W | *Wc* | White | PI587144 |
| KY228 | *Wc* | White | PI587136 |
| Tzi8, Nigeria | *Wc* | White | PI506246 |
| Chile 301, Santiago Chile | *Wc* | White | PI485410 |
| Santander S 356, Columbia | *Wc* | White | PI445401 |
| Boyaca 462, Columbia | *Wc* | White | PI444165 |
| NC336 | *Wc* | White | Ames 27164 |
| CML10 | *Wc* | White | Ames 27072 |
| K55, Kansas, USA | *Wc* | White | Ames 22754 |
| Mexico 37 | *Wc* | White | Ames 19558 |
| White Dent OP | *Wc* | White | Ames 04836 |
| Hays White, WI, USA | *Wc* | White | Ames 01829 |
| Cuzco 9, Lima Peru | *Wc-f* | pale yellow | PI503671 |
| Huancavelica 147, Peru | *Wc-f* | White | PI571793 |
| CML218 | *Wc-f* | White | Ames 27086 |
| CML91 | *Wc-f* | White | Ames 27079 |
| CML247 | - | White | PI595541 |
| H105W | - | White | PI587127 |
| MO15W | - | White | PI558518 |
| Aguascalientes 8, Mexico | - | White | PI484401 |
| White midget, USA | - | White | NSL5631 |
| NC33 | - | White | Ames 27139 |
| I29 | - | White | Ames 27115 |
| Guanajuato 36, Mexico | - | White | Ames 19481 |
| Guerrero 3, Mexico | - | White | Ames 19467 |

<sup>1</sup> *Wc* genotype was determined by Southern blots probed with *Ccd1* for presence of an intense 6.1 kb Bam HI fragment. *Wc-f* indicates presence of a faint band consistent with a low-copy number *Wc* allele.
<sup>2</sup> Phenotypes were recorded for seed grown this experiment; seg, segregating.

**Figure 6.**
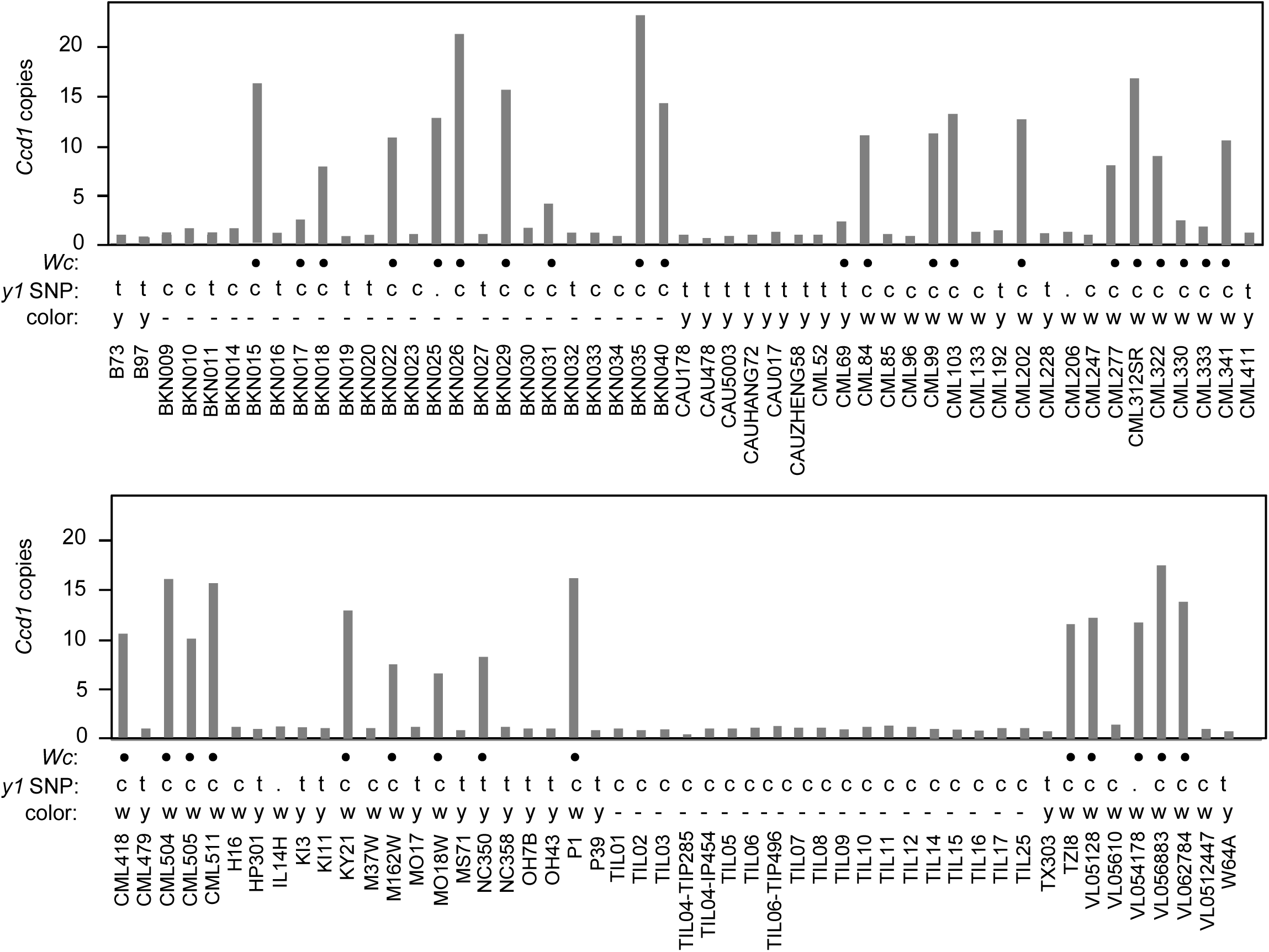
*Ccd1* copy number variation in maize and teosinte populations. K-mer frequency analysis of wgs data (see Materials and Methods) was used to estimated CCD1 gene copy number in genomes of diverse maize landraces, inbreds and teosinte accessions represented in the HapMap2 collection (Chia et al., 2012). *Wc:* •, accessions that contain at least two sequence features that are specific to the *Wc* locus (Table 1). *y1* SNP: homozygosity for T (t) and C (c) variants of PZE0680879922; (.), not determined. Color: grain-color phenotype; y, yellow; w, white; -, data not available.

## Discussion

Our results indicate that a *Ccd1* gene cluster at the *Wc* locus is the basis for a dominant white endosperm phenotype that has likely contributed to human selection for grain color (Fig. 1). Diversity among the *Wc* alleles accounts for extensive *Ccd1* copy-number variation observed in maize. We show that the *Wc* locus was created by macro-transposition and duplication of a chromosome segment containing *Ccd1* and several nearby genes, a process that was mediated by a pair of *Tam3L* transposons. Subsequent tandem duplication of *Ccd1* at the new locus likely set up further expansion and variation of *Ccd1* copy number in *Wc* alleles through unequal crossing-over. Remarkably, transcriptome data indicate that *Ccd1* expression in diverse maize tissues is directly proportional to *Ccd1* copy number over a range of at least 1 to 12 copies per genome. While *Wc* is thus far detected only in maize, its broad geographic distribution is consistent with creation of the locus prior to dispersal of maize from its center of origin in Central Mexico. Interestingly, in diverse landraces as well as modern inbreds, *Wc* is most often found in white-grain varieties that are also homozygous for recessive alleles of *y1* that have little capacity for carotenoid biosynthesis in the endosperm. We suggest that *Wc* contributed to human selection for grain-color by enhancing the *y1* white-endosperm phenotype.

### *Hat* transposons are a potent source of structural diversity in the maize genome

The *Wc* locus illustrates the potency of *Hat* family transposable elements in generating novel structural-genetic variation in the maize genome. Our model (Figs 2 and FigS4) for the creation of *Wc* and amplification of *Ccd1*, builds on previous analyses of the *Ac/Ds* system in maize (Ralston et al., 1989; Zhang et al., 1999, 2013, 2014; Huang and Dooner, 2008). These studies document a variety of chromosome rearrangements arising from interactions between compatible ends of nearby *Ac/Ds* elements. The capacity for macro-transposition in the *Ac/Ds* system is augmented by 1) preferential transposition of *Ac* during DNA replication and 2) the propensity for elements to transpose to nearby sites. Our results indicate that *Tam3L* has similar characteristics. The macro-transposon structure is confirmed by wgs data and PCR. These results together confirm presence of *Tam3La*-right and *Tam3Ld*-left junctions that share a matching 8-bp host-site duplication (Fig 3). The mechanism responsible for tandem duplication of *Ccd1* at *Wc* is less clear. While previously-documented mechanisms for *Ac-*induced DNA re-replication (Zhang et al., 2014) do not account for all aspects of the *Wc* structure, presence of repeats punctuated by the composite *Tam3Lc* sequence implicates *Tam3L* transposons in their formation. Once formed, a partial duplication of the 28-kbp sequence (e.g. Fig S4d) would have enabled expansion of copy number by unequal recombination of *Wc* alleles. Overall, the *Wc* repeats are highly uniform, compatible with a relatively young age. However, variation in *Ccd1:P450* copy-number ratio indicates that *Wc* alleles contain at least two classes of repeats that have diverged through partial or complete loss of *P450* (Fig 4). Hence, the relatively young *Wc* complex will likely continue to evolve toward greater structural heterogeneity as individual repeats diverge in ways that may affect dynamics of recombination.

### Quantitative variation in *Ccd1* expression is proportional to copy-number

Our results indicate that copy-number variation at *Wc* causes proportional quantitative variation in *Ccd1* expression. Remarkably, the gene dosage response is linear up-to at least 12 copies per genome (Fig 5). By contrast, the adjacent gene in the *Wc* repeat, *Tglu*, showed no correlation between expression and gene dosage. Clearly, gene amplification alone is not sufficient to produce a stable, proportional dosage response. While the basis for this intriguing, qualitative difference in dosage dependence of the *Ccd1* and *Tglu* genes is unclear, we speculate that the *Tam3L* insertion at the 5’-end results in more or less constitutive expression of *Ccd1* gene copies. In any case, *Wc* offers a unique opportunity for investigating the effects of tandem duplication on chromatin structure and gene expression.

### Haplotype diversity at *Wc* and selection for grain color

The *Wc* locus most likely originated shortly after the domestication of maize from teosinte, but prior to dispersal of maize from its center of origin in Mexico. In modern maize, the *Wc* locus is broadly distributed in white-grain inbreds and landraces from North, Central and South America as well as Africa (Tables 1 and 2). In contrast, the locus is not detected in any of the 21 teosinte (*Z. mays, parviglumis*) accessions characterized in this study (Fig 6).

The parallel and independent diversification of the multi-copy *Wc* and single-copy *Ccd1r* haplotypes in maize is intriguing because the variation at these loci would potentially support selection for yellow- as well as white-grained maize. On one hand, in a *y1* background, selection for increased *Ccd1* copy-number at *Wc* potentially contributed to breeding of white-endosperm varieties. Conversely, helitron insertions in B73 and OH43 haplotypes that displace or disrupt upstream regulatory sequences of *Ccd1r* would possibly enhance carotenoid accumulation by attenuating carotenoid turnover in yellow-endosperm. The B73 and OH43 variants together show evidence of enrichment in yellow inbreds relative to white inbreds (Χ^2^, p=0.0065).

Although the striking dominant-white phenotype of *Wc* in yellow maize (Y1) was reported a century ago (White, 1917), its contribution to selection and breeding of both traditional and modern white-grain maize has been largely un-appreciated. Homozygous *y1* progeny obtained from crossing white (*y1*) and yellow (*Y1*) inbreds often have off-white ("dingy") phenotypes due to presence of residual pigment (Poneleit, 2001; Stinard, 2010). The off-color phenotype is attributable, at least in part, to *Brown aleurone-1* (*Bn1*, on chromosome 7), which causes accumulation of an unidentified yellow-brown pigment in aleurone (Stinard, 2010). In a *Bn1 y1* background, *Wc* alleles inhibit accumulation of the yellow-brown pigment, thus producing a more intense white-endosperm phenotype (Stinard, 2010; Supplemental Fig S6). The broad-spectrum CCD1 activity could conceivably degrade the product of the *Bn1* pathway.

### Persistence of *Wc* in yellow maize

*Wc* is also occasionally found in yellow-grain maize (*Y1* background). Examples include two modern inbreds, NC350 (8 *Ccd1* copies) and CML69 (2 *Ccd1* copies), as well as landraces from Peru and Central Mexico (Table 2). The majority of these landraces segregate a mixture of white- and yellow-grain phenotypes. Other historically important groups of *Wc Y1* maize include ‘*White Cap Yellow Dent’* and similar open-pollinated varieties that were grown widely in North America in the 19th and early 20th centuries (Brink, 1930). The term "white-cap" was also widely applied to flint landraces grown by Native Americans in Northeastern United States and Eastern Canada (Brink, 1930). While these varieties were not included in our survey, the long history of the “white-cap” phenotype implies that *Wc Y1* genotypes have been utilized through centuries of human cultivation.

The relative rarity of *Wc Y1* in modern inbreds is likely due at least in part to active selection against *Wc* in breeding maize hybrids. Brink (1930) cited two explicit sources of bias against using *Wc Y1* genotypes during formative years of the hybrid seed industry. First, by 1930 *Wc Y1 ‘White Cap Yellow Dent’* varieties were known to have reduced pro-vitaminA content relative to yellow-dent (*wc Y1*) maize affecting their value as livestock feed (Russel, 1930). Second, obtaining a uniform endosperm color in the grain harvested from hybrid plants was an important breeding objective. In promoting utilization and marketing of hybrids, uniform color was highlighted as a contrast with the variation typical of competing, open-pollinated varieties. Achieving uniform color in double-cross hybrids common at that time required that the four inbred parents all be either *Wc* or non-Wc.

Where *Wc* occurs in yellow maize, an increased rate of carotenoid turnover in endosperm is likely to increase production of apo-carotenoid compounds that may contribute to grain quality. Notably, apo-carotenoid products of CCD1 are important determinants of food taste and aroma (Vogel et al., 2008). Some CCD1 products have also been implicated in other biological processes including formation of mycorrhizal symbioses in roots (Sun et al., 2008). The relative importance of aromatic/taste phenotypes of *Wc* would likely depend on how the maize crop was utilized. Peak expression of *Ccd1* during mid grain-fill could lead to production of volatiles with potential to increase quality of kernels harvested early for fresh consumption (e.g. sweet corn or Mexican etole). In New England, the preferred maize for preparation of “johnny cakes” is ‘*Rhode Island White Cap’*, a modern descendant of Native American white cap landraces (Thomas, 1911). Because CCD1 protein is localized to the cytosol (Tan et al., 2003) the enzyme *in vivo* would normally be expected to have limited access to carotenoids that are located primarily in plastid membranes. However, substrate availability would likely increase as cells in the endosperm undergo desiccation during seed maturation, thus accounting for late onset of visible whiteness for the *Wc* phenotype. As a result, the potential for apo-carotenoid production is likely to be comparatively high in freshly-harvested grain. Together these considerations indicate a rich potential for interactions between *Wc* genotypes and the diverse cultural practices and culinary customs built around maize.

The agricultural genomics of *Wc* presented here shows how creation of this unusual locus provided a foundation for human selection of white- and yellow-grain maize. Molecular dissection of the *Wc* locus itself also provides a striking example of transposon remodeling that has altered a genome in a way historically important to humankind.

## Materials and Methods

### Genetic stocks

The reference *Wc Y1* stock, *wc Y1* maintence stock, County White maize lines, and teosinte (Z.m. parviglumis) accessions were obtained from the Maize Coop Genetics Stock Center (Urbana, Illinois). Six diverse inbred stocks were a gift of Joachim Messing, Waksman Institute, Rutgers University. The Silver Queen hybrid sweet corn harboring a dominant white allele was obtained from Dr, L. Curtis Hannah at the University of Florida. Additional accessions listed in Table 2 were obtained from the USDA North Central Regional Plant Introduction Station in Ames, IA. Maize lines were grown at the University of Florida field station in Citra, FL.

### Nucleic Acid Methods

DNA extraction, Southern analysis, sequence determination and other routine molecular biology methods were conducted as previously described (Tan et al., 1997, 2003). The *Ccd1* genomic DNA region was cloned via construction and screening of a lambda phage genomic library prepared from a *Wc wc* heterozygote as previously described (Tan et al., 1997). Briefly, genomic DNA was digested with BamHI and resolved through 10 – 35% sucrose gradient centrifugation at 26,000 ×g for 24 hrs at 4°C. The fraction enriched in 6.5 kb fragments was purified and ligated to lambda-ZAPII vector (Stratagene). The library was screened using a 1.1 kb *Ccd1* partial cDNA (EST CSU453) as a probe. The genomic sequence was extended by PCR amplification of a 2 kb fragment from *Wc* genomic DNA that included the presumptive translation start.

### Cloning of *Ccd1* cDNA and partial genomic clones from maize and teosinte

A near full-length cDNA clone containing the complete coding sequences of *Ccd1* was obtained by RT-PCR with forward primer (5’-CCCTTCGCTACAAGCCTACA) and reverse primer (5’-TTCGAATACACGTCCTGCAA). RNA extracted from developing *Wc* kernels at 18 days after pollination was used to synthesize the cDNA. The PCR products were cloned in pCR4-TOPO vector and completely sequenced. Several genomic fragments identified from Southern analysis were cloned by construction and screening of λ-phage libraries. Briefly, genomic DNA was digested with appropriate DNA restriction enzymes and fractionated via a 15%-40% sucrose gradient centrifuged at 25,000 x g at 4°C for 24 hrs in a swing bucket rotor. Selected DNA fractions were ligated into phage cloning vectors (λ-ZAP II or λ-ZAP Express, Stratagene) and packaged according to the manufacturer’s instruction. The library was screened and positive clones were isolated and converted into phagemid by in vivo excision. The plasmids were sequenced (Genbank accessions genomic clones DQ100348, DQ100347 and *Wc* cDNA DQ100346).

### BAC library construction and sequencing

A custom bac library was constructed from genomic DNA of the homozygous *Wc* reference stock and screened with a *Ccd1* cDNA probe by the Bio S&T Inc. (Montreal, Canada). Two positive clones (F19 and H10) were identified. The BAC clones were characterized by restriction mapping and partial sequencing of selected subcloned restriction fragments. Of the two clones, H10 extended farthest into the *Wc* locus, and was selected for complete sequencing and assembly using a combination of circular-concensus and linear long format reads from the PacBiosystems instrument. Trimmed circular-concensus sequence reads (>5,000 bp) were assembled into contigs using CAP3 (Huang and Madan, 1999) and linear long reads were assembled using CANU (Koren et al., 2016). The contigs were further evaluated and assembled manually to obtain an assembly (Genbank accession KX760165) that was consistent with the bac restriction map, bac end sequences, wgs analysis, and subclone sequences.

### Quantitative real time RT-PCR

Total RNA was extracted using RNeasy (Qiagen, Germany) and treated with RNase-free DNase. The complete removal of DNA was verified by a quantitative real time RT-PCR analysis without reverse transcription. The conditions used are as described in detail previously (Tan et al., 2003). For TacMan qPCR primers used for *Ccd1* were forward (5’-GGGAAGAGGGTGATGAAGTTGT) and reverse (5’-TGATATCCATTCACCTTGTCCAAA), and the probe was 5’-CTCATTACCTGCCGCCTTGAGAATCCA. The probe was labeled with fluorescent reporter dye 6-carboxyfluorescein (FAM) at 5’ and 6-carboxy-N,N,N ’,N’-tetramethylrhodamine (TAMRA) at 3’. The standard curve was derived with a plasmid containing *Ccd1* cDNA. Reactions were carried out in the GeneAmp 5700 Sequence Detection (Perkin-Elmer). The transcript abundance was normalized as copy number per nanogram of total RNA. qPCR of pericarp tissue was performed as described by Sun et al. (2008) using 5’-CTGCTGTGGATTTTCCTCGTG and 5’-TATGATGCCAGTCACCTTCGC as forward and reverse primers, respectively. Relative expression levels were calculated from E^-ΔCt^ values setting the mean of the B73 wc control to 1.

### Identification of the *Tam3Ld* left junction sequence

To identify the left border sequence of the *Wc* macro-transposon, we employed a modified TAIL-PCR protocol which used four AD3 primers (AD3-1 to 4), each of which contains an AD3 primer fused with an arbitrary degenerate primer (Liu and Chen 2007). Three nested primers based on the predicted *Tam3Ld* flanking sequence (Wc3P-R3, Wc3P-R4 and Wc3P-R5) were used with the AD3 primers. Following three rounds of TAIL-PCR fragments were sequenced and analyzed to identify products that contained *Tam3L-*left termini. The sequences were in turn used to design a *Tam3L*-specific primer (Wc3P-R5) which was then used with Wc3P-R3 to amplify the left border of the *Tam3Ld* candidate sequence. The resulting *Tam3Ld* left flanking sequence was contained an 8-bp target site duplication (GTTCTAGT) that matched the right junction of *Tam3La* confirming the macro-transposon hypothesis.

**Table 3.**
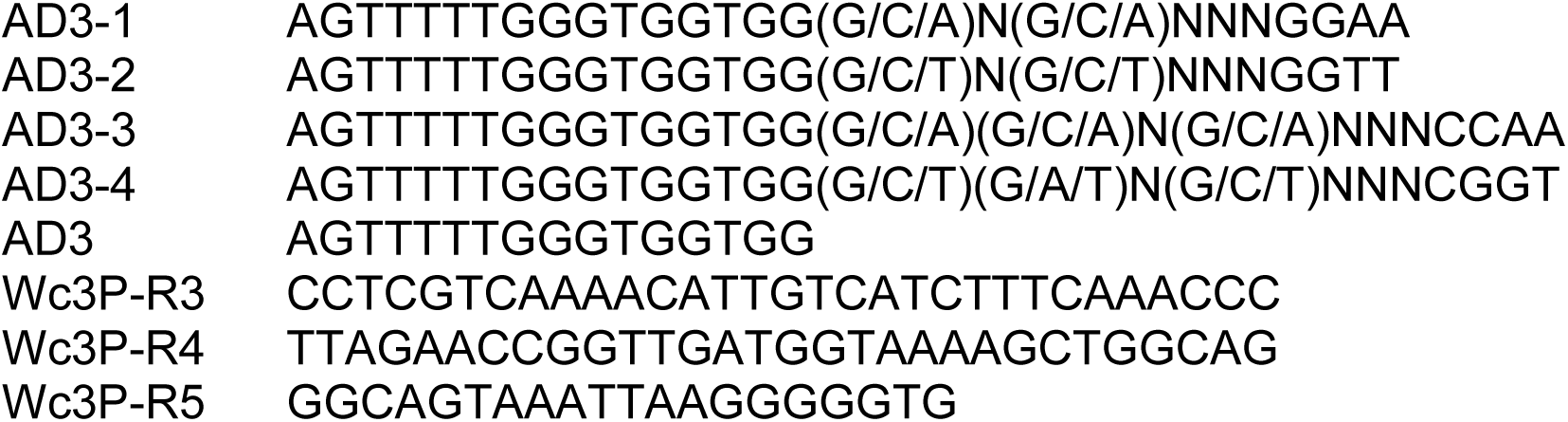
Primers used to identify the Tam3Ld left junction

### Quantification of *Ccd1* copy number in the *Wc* reference allele

Real-time quantitative PCR was used to determine the gene copy number of *Ccd1* in each DNA sample. A single copy gene *Vp14* (Tan et al., 1997) was used as an internal standard. In addition, the inbred line B73 which was confirmed to contain single copy of *Ccd1* by hybridization and sequencing was used as a standard to normalize the *Ccd1* probe. To increase the accuracy of real-time quantitative PCR, genomic DNA samples were digested with EcoRI to completion, generating *Vp14* and *Ccd1* fragments in a 6 to 7 kb size range. The DNA’s were further purified by a Turbo genomic DNA purification kit (Q-Biogene). The concentration of the DNA’s was determined spectrophotometrically. Equal amounts of DNA were analyzed by real-time quantitative PCR. The analysis conditions were same as the real-time quantitative RT-PCR described above except without reverse transcription. *Ccd1* primers and probe were same as above. The *Vp14* primers are forward (5’- GCTGGCTTGGCTTGTATACTCTG-T) and reverse (5’-CCATCAGTCATATACTGTGAACAAATGT), the gene specific probe is (5’-CACGCACCGATAGCCACAGGGAA) labeled with FAM and TAMRA at 5’ and 3’ respectively.

### Copy number estimation in maize genomes by analysis of k-mer frequencies

Frequencies of 22-mers in the B73 reference genome were profiled using JELLYFISH (Marçais and Kingsford, 2011). The resulting database was then queried with 22-mers from 39,424 genes in the maize filtered gene set (gramene.org) to identify a subset of genic 22-mers that were single-copy in the B73 genome. Frequencies of the resulting set of 124 million single-copy, genic 22-mers were in turn profiled in wgs sequence data obtained from the SRA archive (ncbi.nlm.nih.gov) for each of the 102 maize and teosinte accessions in the HapMap2 collection (Chia et al., 2012). Gene copy numbers in each genome were then estimated by normalizing the average frequency of single-copy 22-mers from *Ccd1r* to the average frequency of 124 M genic single-copy 22-mers in wgs data for each inbred. The estimated effective sequence coverage of each genome is listed in Supplemental Table S1.

### Analysis of *Wc* and *Ccd1r* allele-specific features in maize genomes

The wgs data from HapMap2 genomes was searched for sequence reads that contained diagnostic features of the *Wc* locus and *Ccd1r* alleles using the GREP utility. Simple text searches were made in both orientations using 18-22 base sequences that were unique to transposon insertion sites and other characteristic features of *Wc* or *Ccd1r* alleles. Sequence reads identified by text searches were then validated by full-length blastn alignment to the *Wc* bac assembly and B73 reference genome (Schnable et al., 2009) sequences.

## Acknowledgements

We thank Dr. Daniel Ngu and Dr. Patrick Schable (Iowa State University) for permission to use NAM inbred RNAseq data deposited at QTELLER.ORG. We are grateful to Phil Stinard and Marty Sachs at the Maize Coop Genetics Stock Center for drawing our attention to the interaction between *Wc* and *Bn1*, stimulating discussions, permission to cite Maize Genetics Newsletter notes, and provision of genetic stocks. This work was supported by grants to DRM and KEK from the National Science Foundation (IOS:1116561) and USDA-NIFA (2011-67003-30215).

## References

Brink, R. A. Some problems in the utilization of inbred strains of corn (Zea mays). The American Naturalist 64, 525–539 (1930).

Buckner, B., Kelson, T.L. & Robertson, D.S. Cloning of the *y1* locus of maize, a gene involved in the biosynthesis of carotenoids. Plant Cell 2, 867–876 (1990).

Buckner, B., Miguel, P.S., Janick-Buckner, D. & Bennetzen, J.L. The *Y1* gene of maize codes for phytoene synthase. Genetics 143, 479–488 (1996).

Chia, J.-M., et al. Maize HapMap2 identifies extant variation from a genome in flux. Nat. Genet. 44, 803–807 (2012).

DeBolt, S. Copy number variation shapes genome diversity in Arabidopsis over immediate family generational scales. Genome Biol. Evol. 2, 441–453 (2010).

Fu, H. & Dooner H. K. Intraspecific violation of genetic colinearity and its implications in maize. Proc. Natl. Acad. Sci. USA 99, 9573–9578 (2002).

Giguere, V., Ong, E.S., Segui, P., & Evans, R.M. Identification of a receptor for the morphogen retinoic acid. Nature. 330, 624–629 (1987).

Gomez-Roldan, V. et al. Strigolactone inhibition of shoot branching. Nature 455, 189–194 (2008).

Han, F., Lamb, J.C. Yu, W., Gao, Z. & Birchler, J.A. Centromere function and nondisjunction are independent components of the maize B chromosome accumulation mechanism. Plant Cell 19, 524–533 (2007).

Hannah, L.C. & McCarty, D.R. Maize Genetics Coop Newsletter 65, 62 (1991).

Hardigan, M.A. et al. Genome reduction uncovers a large dispensable genome and adaptive role for copy number variation in asexually propagated *Solanum tuberosum*. Plant Cell 28, 388–405 (2016).

Huang, J.T. & Dooner, H.K. Macrotransposition and other complex chromosomal restructuring in maize by closely linked transposons in direct orientation. Plant Cell 20, 2019–2032 (2008).

Huang, X. & Madan, A. CAP3: A DNA sequence assembly program. Genome Res. 9, 868–877 (1999).

Kempken, F. & Windhofer, F. The hAT family: a versatile transposon group common to plants, fungi, animals, and man. Chromosoma 110, 1–9 (2001).

Koren, S., Walenz, B.P., Berlin, K., Miller, J.R. & Phillippy, A.M. Canu: scalable and accurate long-read assembly via adaptive k-mer weighting and repeat separation. bioRxiv. p.071282 (2016).

Liu, Y.G. & Chen, Y. High-efficiency thermal asymmetric interlaced PCR for amplification of unknown flanking sequences. Biotechniques 43, 649–656 (2007).

Marçais, G., & Kingsford, C. A fast, lock-free approach for efficient parallel counting of occurrences of k-mers. Bioinformatics 27, 764–770 (2011).

Palaisa, K.A., Morgante, M., Williams, M. & Rafalski, A. Contrasting effects of selection on sequence diversity and linkage disequilibrium at two phytoene synthase loci. Plant Cell 15, 1795–1806 (2003).

Palaisa, K., Morgante, M., Tingey, S. & Rafalski, A. Long-range patterns of diversity and linkage disequilibrium surrounding the maize *Y1* gene are indicative of an asymmetric selective sweep. Proc. Natl. Acad. Sci. USA 101, 9885–9890 (2004).

Poneleit, C.G. Breeding white endosperm corn. In Specialty Corns (2nd edition). Ed. A. R. Hallauer. CRC Press. Boca Raton, FL. (2001).

Ralston, E., English J. & Dooner, H.K. Chromosome-breaking structure in maize involving a fractured *Ac* element. Proc. Natl. Acad. Sci. USA 86, 9451–9455 (1989).

Russel, W.C. The vitamin A content of yellow and white-capped yellow dent corn. J. Nutrition 2, 265–268 (1930).

Schnable, P.S., et al. The B73 maize genome: complexity, diversity, and dynamics. Science 326, 1112–1115 (2009).

Schwartz, S.H., Tan, B.C., Gage, D.A., Zeevaart, J.A., & McCarty, D.R. Specific oxidative cleavage of carotenoids by VP14 of maize. Science 276, 1872–1874 (1997).

Springer, N.M. et al. Maize inbreds exhibit high levels of copy number variation (CNV) and presence/absence variation (PAV) in genome content. PLoS Genet. 5, e1000734 (2009).

Sun, Z. et al. Cloning and characterisation of a maize carotenoid cleavage dioxygenase (ZmCCD1) and its involvement in the biosynthesis of apocarotenoids with various roles in mutualistic and parasitic interactions. Planta 228, 789–801 (2008).

Stinard, P. S. (2010) Isolation and characterization of a dominant inhibitor of Bn1. Maize Genetics Cooperation Newsletter 84:42–43.

Studer, A., Zhao, Q., Ross-Ibarra, J. & Doebley, J. Identification of a functional transposon insertion in the maize domestication gene *tb1*. Nat. Genet. 43, 1160–1163 (2011).

Tan, B.C. et al. Molecular characterization of the Arabidopsis 9-*cis* epoxycarotenoid dioxygenase gene family. Plant J. 35, 44–56 (2003).

Tan, B.C., Schwartz, S.H., Zeevaart, J.A., & McCarty, D.R. Genetic control of abscisic acid biosynthesis in maize. Proc. Natl. Acad. Sci. USA 94, 12235–12240 (1997).

Tracy, W. F. Sweet corn. in Specialty Corns Second Edition. ed. A. R. Hallauer. CRC, Boca Raton, FL. pp.155–199 (2000).

Thomas, E.K. Corn and its uses. J. Education. 74, 327 (1911).

Umehara, M. et al. Inhibition of shoot branching by new terpenoid plant hormones. Nature 455, 195–200 (2008).

Vogel, J.T., Tan, B.C., McCarty, D.R. & Klee, H.J. The carotenoid cleavage dioxygenase 1 enzyme has broad substrate specificity, cleaving multiple carotenoids at two different bond positions. J. Biol. Chem. 283, 11364–11373 (2008).

White, O.E. Inheritance of endosperm color in maize. Amer. J. Bot. 4, 396–406 (1917).

Yu, J.M., Holland, J.B., McMullen, M.D., & Buckler, E.S. Genetic design and statistical power of nested association mapping in maize. Genetics 178, 539–551 (2008).

Zeevaart, J.A., Heath, T.G. & Gage, D.A. Evidence for a universal pathway of abscisic acid biosynthesis in higher plants from o incorporation patterns. Plant Physiol. 91, 1594–1601 (1989).

Zhang, J. & Peterson, T. Genome rearrangements by nonlinear transposons in maize. Genetics 153, 1403–1410 (1999).

Zhang, J., Zuo, T. & Peterson, T. Generation of tandem direct duplications by reversed-ends transposition of maize ac elements. PLoS Genet. 9, e1003691 (2013).

Zhang, J., Zuo, T., Wang, D. & Peterson, T. Transposition-mediated DNA re-replication in maize. Elife 3, e03724 (2014).

Zhu, C. et al. Combinatorial genetic transformation generates a library of metabolic phenotypes for the carotenoid pathway in maize. Proc. Natl. Acad. Sci. USA 105, 18232–18237 (2008).

